# Fast, multicolor 3-D imaging of brain organoids with a new single-objective two-photon virtual light-sheet microscope

**DOI:** 10.1101/461335

**Authors:** Irina Rakotoson, Brigitte Delhomme, Philippe Djian, Andreas Deeg, Maia Brunstein, Christian Seebacher, Rainer Uhl, Clément Ricard, Martin Oheim

## Abstract

Human inducible pluripotent stem cells (hiPSCs) hold a large potential for disease modeling. hiPSC-derived human astrocyte and neuronal cultures permit investigations of neural signaling pathways with subcellular resolution. Combinatorial cultures, and three-dimensional (3-D) embryonic bodies enlarge the scope of investigations to multi-cellular phenomena. A the highest level of complexity, brain organoids that – in many aspects – recapitulate anatomical and functional features of the developing brain permit the study of developmental and morphological aspects of human disease. An ideal microscope for 3-D tissue imaging at these different scales would combine features from both confocal laser-scanning and light-sheet microscopes: a micrometric optical sectioning capacity and sub-micrometric spatial resolution, a large field of view and high frame rate, and a low degree of invasiveness, i.e., ideally, a better photon efficiency than that of a confocal microscope. In the present work, we describe such an instrument that belongs to the class of two-photon (2P) light-sheet microsocpes. Its particularity is that – unlike existing two- or three-lens designs – it is using a single, low-magnification, high-numerical aperture objective for the generation and scanning of a virtual light sheet. The microscope builds on a modified Nipkow-Petráň spinning-disk scheme for achieving wide-field excitation. However, unlike the common Yokogawa design that uses a tandem disk, our concept combines micro lenses, dichroic mirrors and detection pinholes on a single disk. This design, advantageous for 2P excitation circumvents problems arising with the tandem disk from the large wavelength-difference between the infrared excitation light and visible fluorescence. 2P fluorescence excited in by the light sheet is collected by the same objective and imaged onto a fast sCMOS camera. We demonstrate three-dimensional imaging of TO-PRO3-stained embryonic bodies and of brain organoids, under control conditions and after rapid (partial) transparisation with triethanolamine and /ormamide (RTF) and compare the performance of our instrument to that of a confocal microscope having a similar numerical aperture. 2P-virtual light-sheet microscopy permits one order of magnitude faster imaging, affords less photobleaching and permits better depth penetration than a confocal microscope with similar spatial resolution.

## INTRODUCTION

The development of pharmacological treatments for neuropsychiatric and neurodegenerative diseases has been hampered by the poor availability of preclinical models that adequately capture the complexity of human disorders [2].

Human inducible pluripotent stem cells (hIPSCs) offer a promising platform for disease modeling and drug screening. A comparably new technique is the directed differentiation and reprogramming of patient fibroblasts into neurons, astrocytes, microglia and oligodendrocytes. Their combinational culture permits the growth of embryonic bodies (EBs) and brain organoids, 3-D cultures that – in many aspects – recapitulate the development of the human brain [3; 4]. Together, hIPSCs, EBs, and brain organoids enable observations and experiments that were previously inconceivable, neither on human subjects, nor in animal models [5; 6]. Recent reports of functional, fully vascularized brain organoids have spurred hopes of growing even larger 3-D cell assemblies [7], bringing the hitherto theoretical *‘brain in a vat’*^1^ within reach of the imaginable.

Elucidation of neural circuit (dys-)function would benefit from the detailed, 3-D visualization of the fine structure of neurons, astrocytes and blood vessels over large fields of view and deep in tissue. Large-scale neuroanatomical imaging has become possible in cleared tissue sections [8], brain organoids [9] or even entire brains [10], but in many cases the resolution is rather at level of cell bodies that at the synaptic scale. In addition to the difficulties associated with transparisation and tissue shrinking, imaging of large tissue volumes at spatial high-resolution presents considerable challenges: confocal and two-photon (2P) laser scanning microscopies set the ‘gold-standard’ for diffraction-limited fluorescence imaging, but – being in most of their implementations point-scanning, i.e., sequential techniques – the image acquisition is often painstakingly slow. Particularly, the reconstruction of large volumes often requires hours if not days of recording, putting high demands on mechanical stability of the microscope, photostablity of the used fluorescent dyes, and incurring considerable cost for beam time. Line- and multi-spot scanning schemes overcome these limitations by parallelizing the *excitation,* but they trade off resolution against speed and they often have relatively small fields of view, requiring image stitching for larger fields.

On the other end, selective-plane illumination microscopes (SPIM) [11] or light-sheet microscopes [12] decouple fluorescence excitation and collection by using orthogonal illumination and detection optical paths. Light-sheet microscopes have established themselves as efficient workhorses for volume imaging in cleared tissue. It is the parallelization of *both excitation* and fluorescence *detection* that allows for rapid 3-D imaging on these instruments [13; 14]. However, one consequence of the lower-NA illumination and a result of excitation-light scattering in not perfectly transparent samples, is that the axial resolution of light-sheet microscopes remains poor compared to the optical sectioning achieved by spot-scanning microscopes. Improvement has been made with 2P light-sheet excitation [15; 16], by combining 2P-line excitation and confocal slit detection [17], by the use of Airy- [18] or Bessel-beams for excitation [19; 20; 21], or a combination of these techniques [22; 23]. However, many of these recent techniques are not yet commercial and they afford considerable cost and complexity compared to standard 1- and 2P-laser scanning microscopes.

Another limitation of light sheet microscopes results from their orthogonal arrangement of excitation and collection objectives: the need for non-standard procedures for embedding and holding the sample. Variants of light-sheet microscopes in which both illumination and detection objectives are mounted at an oblique angle with respect to the tissue surface and the sample half space is left free exist [24; 25; 26], but they have remained comparably marginal.

An ideal microscope for volume imaging in cleared brain tissue [27] would combine speed, a sub-μm lateral and μm-axial resolution, a mm-field of view, an excitation depth of a few mm, a certain robustness to imperfect sample transparisation and a large free space under the objective.

Here, we present a microscope with excitation- and detection-parallelization that gets close to this ideal by combining advantages of 2P laser-scanning and light-sheet techniques. Our <u>O</u>n-<u>a</u>xis 2-photon light-sheet generation *in-vivo* imaging system (OASIS) uses a vast array of micro lenses arranged in four nested spirals on a single spinning disk to simultaneously scan ∼40 independent excitation spots in the focal plane of a single, long-working distance, low-magnification, high-NA objective [28]. Rotation of the disk at 5,000 rpm results in rapid multi-spot scanning and creates a virtual light sheet in the focal plane of the objective. The fluorescence generated in each of the excitation spots is imaged through the same objective onto a pinhole in the center of each micro lens. With the remainder of the lens made opaque to (scattered) fluorescence by a dichroic coating sparing only the tiny pinhole, only fluorescence emanating from the focus is detected. Each pinhole is imaged onto a large-format scientific Complementary Metal Oxide Semiconductor (sCMOS) camera, allowing near-diffraction limited imaging over a large field of view. This patented optical design, combining micro-lenses and perforated dichroic mirrors on a single-spinning disk, allowed us to retain the in-line, single-objective geometry of a classical microscope without the requirement for orthogonal illumination. As a consequence, our OASIS microscope is more versatile than two- or three-objective light-sheet microscopes. With its compact footprint (43 cm by 12 cm, or 17” × 5”), it can accommodate large samples (cells, slices, explants and entire animals, *in vivo*) without requiring tedious mounting procedures or special sample holders. The OASIS concept combines the optical sectioning, spatial resolution and field-of-view of a 2P-scanning microscope with the speed of a light-sheet microscope. Due to 2P excitation, out-of-focus fluorescence excitation, photo-bleaching and photo-damage are much reduced compared to a classical confocal microscope. We here describe this new microscope and compare it to a confocal laser-scanning microscope (CLSM) for imaging clarified brain organoids with nuclear staining.

## RESULTS AND DISCUSSION

### Wide-field two-photon microscopy at diffraction-limited resolution

With our <u>O</u>n-<u>a</u>xis 2-photon light-<u>s</u>heet generation *in-vivo* imaging system (OASIS), we retain the in-line geometry of a classical upright microscope with a single objective lens. We introduce a novel spinning-disk concept, rethought and specifically designed for wide-field 2P microscopy, **Fig. 1*A***. Briefly, the expanded and shaped beam of a fs-pulsed infrared laser is focused by an array of micro lenses to produce some 40 evenly lit excitation spots. These spots are imaged by the tube lens and objective into the sample plane where they are each spaced, on average, by 28 μm, **Fig. 1*A***, *inset* ➊. A total of almost 5,000 micro lenses are arranged in four nested spirals that scan these spots upon rotation of the disk. At 5,000 rpm (i.e., one turn every 12 ms, much shorter than camera integration times used here) the multispot scanning generates a virtual light sheet permitting wide-field, direct-view 2P imaging. Unlike earlier 2P-spinning disk microscopes [29; 30; 31] that were based on modified Yokogawa-type spinning-disk confocal microscopes, we use a different disk geometry. Where the Yokogawa dual-disk design requires two disks, one with micro lenses, the other one with confocal pinholes, the OASIS microscope relies on a single disk on which micro lenses and pinholes that are arranged on the different faces of the same optical element. Specifically, we do not use an extra dichroic, but each micro lens comes with its own dichroic mirror; the dielectric coating is omitted over a central circular aperture of 60-μm diameter. The use of a single disk is advantageous in view of the large wavelength difference between the near infrared excitation and the visible fluorescence, which had previously reduced the efficiency of 2P-spinning disk miroscopes. However, this simplification comes at a price as it required a complete re-design of the microscope optical path. On the excitation side, the micro lenses must be uniformly illuminated with a collimated fs-pulsed IR laser beam to generate an array of focused spots. On the collection side, to pass the pinholes, the detected fluorescence must arrive focused at the level of the pinholes. The required optical path-length difference between excitation and emission light is achieved by introducing a corrective distance element in the non-infinity space: while the longer-wavelength excitation light passes straight through the device, the shorter-wavelength fluorescence takes a detour and travels a longer path to produce the desired focal offset, *inset* ➋. This patented optical scheme critically relies extremely flat shallow-incidence long-pass dichroic mirrors to preserve the phase front of the beam and maintain the optical resolution.

**Fig. 1.**
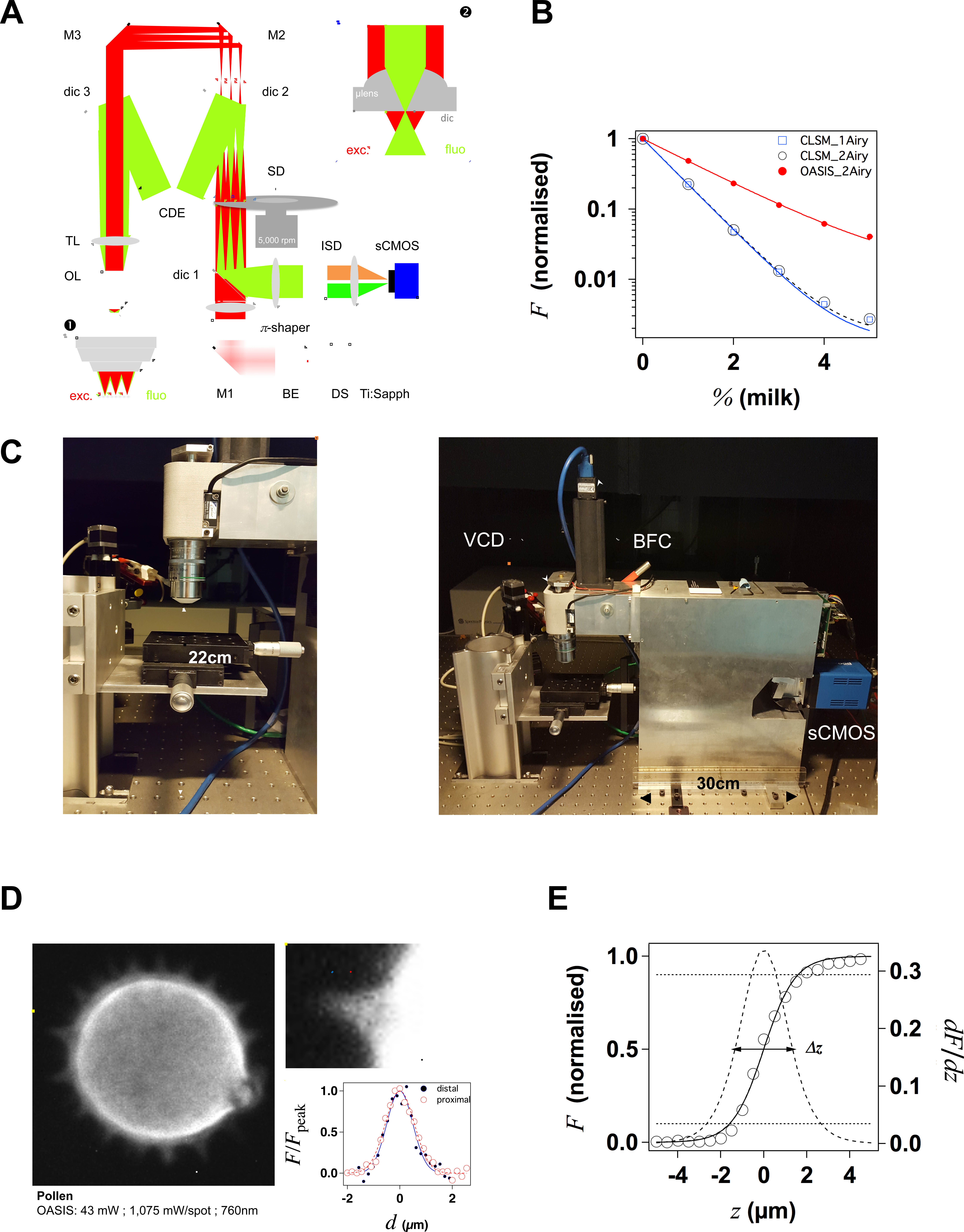
Layout and performance of the OASIS two-photon microscope. **A.** Simplified optical path of the custom 2P virtual light-sheet microscope. *Red:* IR excitation, *green:* fluorescence. Ti:Sapph – fs-pulsed IR laser; DS – DeepSee; BE – beam expander; π-shaper – optical element that converts the Gaussian beam into a top hat profile; M – mirrors; dic – dichroic mirrors; SD – spinning disk; TL – tube lens; CDE – corrective distance element; OL – objective lens; ISD – image-splitting device; sCMOS – camera. *Inset* ➊: generated multi-spot excitation pattern und epi-collection of the generated fluorescence through the same objective. *Inset* ➋: detail of microlens/pinhole/dichroic coating arrangement. Note the offset between the excitation (exc.) and fluorescence foci (fluo.) at the level of the disk, produced by the CDE. **B**. Depth penetration in turbid samples. Log-plot of 2P-excited fluorescence from a green-fluorescent Chroma test slide, topped either with water (0%) or increasing concentrations of milk (a model for the multi-scale scatterers present in tissue), for the OASIS (red dots) and a ZEISS LSM710. Confocal pinhole diameters were 1 and 2 Airy, as indicated. **C**. OASIS prototype, note the compact size and space available around the objective, *inset.* VCD – voice coil z drive; BFC – bright-field camera. **D**. OASIS lateral resolution. Equatorial section through an autofluorescent thorny pollen grain. Scale bar, 10 μm. *Inset:* magnified view of an individual spine and intensity profiles across the lines shown; Scale bar, 2 μm. **E**. OASIS *z*-resolution. Axial-intensity profile measured from a *z*-stack of images acquired from a green fluorescent Chroma slide (solid line), and its derivative *dF/dz* (dashed). The FWHM, equiva-lent to the 10-90% intensity range, was taken as axial optical sectioning capacity *Δz*.

The 2P-excited fluorescence generated in each of the spots is collected through the same objective and imaged onto the tiny pinholes in the dichroic coating of the microlenss (**Fig. 1*A***, *inset* ➋). Relay optics then images these pinholes onto a large-format sCMOS camera. The resulting pixel size in the sample plane is 182 nm, within the Nyquist limit.

This detection is partially confocal (the 60-μm pinhole diameter correspond to a confocal aperture of 2 Airy units), so that only ballistic and snake-like photons but not scattered fluorescence contribute to the signal, as illustrated by the exponential fluorescence drop with increasing sample turbidity. Yet, as a result of the longer wavelength of excitation light, the signal drop observed with the OASIS microscope was only half of that observed with a 1-P CLSM at 633-nm excitation, **Fig. 1*B***. We can attribute the improved depth penetration of the OASIS uniquely to excitation effects, because stopping down the confocal pinhole from 2 to 1 Airy units did not measurably alter the fluorescence decay on the CLSM.

With its small footprint (43 cm by 12 cm) and 22-cm clearance under the objective, the OASIS microscope offers facile access and ample space around the objective, making it an ideal platform for imaging large samples, but also for placing electrodes, application pipettes or external fibers for photostimulation or photochemical uncaging, **Fig. 1*C***. Also, as a re-sult of the large chip format of the sCMOS detector, we could implement simultaneous dual-color detection by way of a custom image splitter, **Fig. S*1***.

With the ×4.6 beam expander and a ×25/NA1.1w low-mag high numerical aperture (NA) dipping objective [28], the OASIS microscope features a field of view with a 200 μm image diagonal. A 5- to 6-fold larger field, making full use of the nominal field-of-view of the same objective would be possible, but it requires more micro-lenses to be illuminated, which in turn requires a more powerful laser than ours.

As a wide-field imaging system, the OASIS microscope simultaneously offers a fast frame rate, a large field of view and it resolves tiny subcellular detail. We illustrate the submicrometric resolution by imaging the fine tip of a spine from an autofluorescent pollen grain, a typical test sample for 2P-microscopes, **Fig. 1*D***.

For estimating the optical sectioning capability of our OASIS microscope, we recorded the axial (*z*-) intensity profile from a green fluorescent Chroma test slide and we quantified the *z*-resolution (Δz) by the full-width at half maximum (FWHM) of a Gaussian fitted with the derivative of the *z*-profile [32], **Fig. 1*E***. Repeating the same experiment on a ZEISS LSM710 demosntrates that OASIS offered a 1.3-fold better optical sectioning than the confocal laser-scanning microscope (with the pinhole diameter set to 2 Airy diameter and with a dipping objective having a similar NA, 1.1w vs. 1.0w; Δz_OASIS_ = 2.75 ± 0.02 μm vs. Δz_CLSM,2Airy_ = 3.47 ± 0.02 μm). In fact, its optical sectioning is close to that of a CLSM with the pinhole stopped down to 1 Airy diameter (Δ*z*_CLSM, 1Airy_ = 2.55 ± 0.02 μm), **Fig. S2**.

The acquisition speed will be in practice be limited by the available signal, but the theoretical minimal exposure time is bounded by the Nyquist limit, i.e., the time required for two full rotations of the spiral on the disk. With a rotation time of 12 ms at 5,000 rpm and four nested spirals, the minimal theoretical exposure time is 3 ms, i.e., 6 ms when taking into account Nyquist’s sampling theorem. Integration times should be multiples of 6 ms for obtaining a homogeneously lit field of view. With the 100-fs pulses and mW average laser power, *<P>,* per illumination spot used here, typical exposure times were of the order of hundred ms, full frame, more than one order of magnitude faster than typical time required a similarly resolved image with a 2P-scanning microscope.

Taken together, our OASIS virtual 2P light-sheet microscope offers better depth penetration, a similar if not better spatial resolution and a considerably higher speed than a conventional CSLM.

### RTF-cleared TO-PRO3-labeled embryos as a test sample for 1- and 2P microscopies

Next, we evaluated the performance of the OASIS microscope for 3-D imaging of partially cleared brain tissue. We sought for a stereotypic, sparsely but homogenously labeled and thick sample. To allow for a direct comparison between 1P-CLSM and our 2P virtual-light sheet microscope, this labeling needed to be suitable for both linear- and non-linear excitation. To minimize scattering and improve the depth penetration, we searched for a red-exciting, deep-red emitting fluorophore. With these constraints in mind we opted for the nuclear stains TO-PRO3 and Methyl Green (MG), having 1P-fluorescence excitation/emission maxima of 642/661 nm [33]) and 632/650 nm [34], respectively.

During preliminary experiments in 7-μm thin sections of a fixed (E14.5) mouse embryo, we found TO-PRO3 fluorescence to be 2.1-fold brighter than that of MG upon 633-nm excitation. We observed an even larger intensity ratio (×5.5) upon 2P-excitation at 760 nm, **Fig. 2*A***. Although non-linear excitation of TO-PRO3 at 1,100 nm has been reported [35; 36] we measured the 2P-action spectra and found peak excitations at 760 and 750 nm for TO-PRO3 and MG, respectively, **Fig. 2*B***. Compared to the reported 1,100-nm excitation these shorter wavelengths are within the tuning range of the standard Ti:Saphh laser and they minimize thermal damage from near-infrared absorption and focal heating [37], a particular concern for the multi-spot excitation scheme used here.

**Fig. 2.**
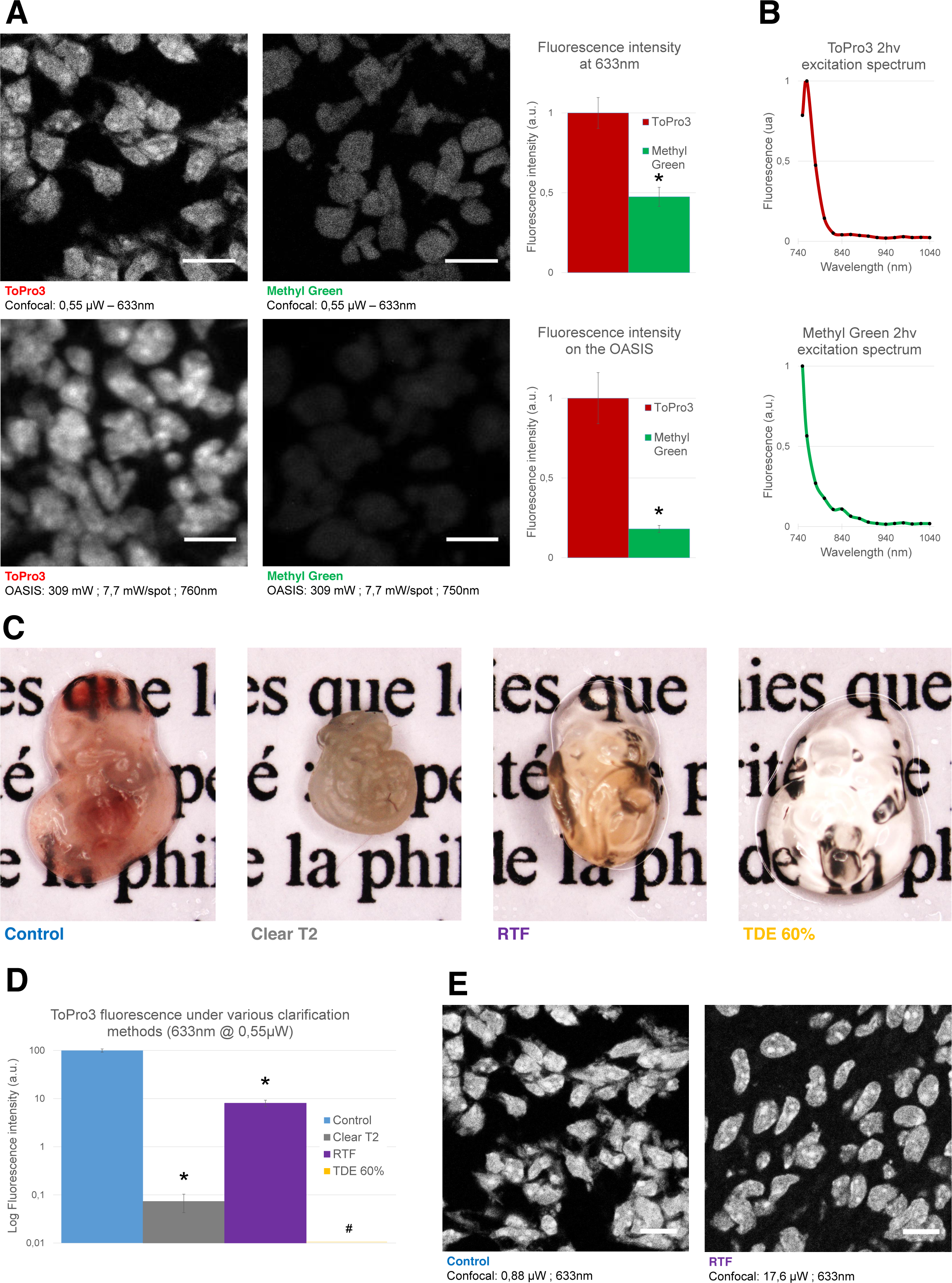
Comparison of the used nuclear stains and clearing methods. **A.** *Left,* optical section of a slice from an E14.5 embryo, labeled with TO-PRO3 or Methyl Green and observed on a confocal laser scanning microscope (CLSM, top) or on the OASIS microscope (*bottom*). Scale-bar, 10 μm. *Right,* relative fluorescence intensity of TO-PRO3 and Methyl Green in nuclei of E14.5 embryo slices measured on a CLSM (*top right*) or on the OASIS (*bottom right).* **B.** TO-PRO3 (*top*) and Methyl Green (*bottom*) 2P-excitation spectra. Color code as in (A). **C.** Macrophotographies of E10.5 embryos in control (PBS) and following three clearing protocols (Clear^T2^, RTF and TDE 60%). Note the variable degree of transparisation and the volume change. **D.** Relative fluorescence of TO-PRO3-positiev nuclei in E14.5 embryo slices observed on a CLSM after the same clearing protocols as above. Note the log-scale. * = *P* < 0.0001. **E.** Fluorescence loss upon clearing requires high laser powers. Confocal micrographs of slices from an E14.5 embryo labeled with TO-PRO3 under control (*left*) and after RTF clearing (*right*) along with the laser powers required to attain the same signal-to-background ratio. Scale-bar, 10μm.

We next optimized the tissue transparisation procedure. Among the available methods (see [38] for review), we focused on TDE [39], Clear^T2^ [40] and RTF clearing [41]. The rationale was that these methods require only short clearing episodes and they use solvents compatible with dipping objectives. Mouse embryos were most transparent with TDE (60%), followed by RTF and, by far, Clear^T2^, for which the tissue was even more opaque than the non-cleared control (probably due to volume shrinkage), **Fig. 2*C***.

In clearing, transparency is one issue, fluorescence preservation another. Depending on the very method used, the observed signal loss was dramatic, with a 99.9% and 92% attenuation of TO-PRO3 fluorescence following Clear^T2^ and RTF clearing, respectively, **Fig. 2*D***. Increasing the laser power by a factor of 20 allowed us to acquire confocal images of TO-PRO3 stained nuclei in slices of RTF-cleared embryos, **Fig. 2*E***, whereas TDE clearing attenuated the fluorescence to undetectable levels, **Fig. S*3***. To develop an order-of-magnitude idea of the laser powers required for obtaining similar signal-to-noise levels with the OASIS and CSLM, we finally compared images acquired upon 1-(at 633 nm) and 2-excitation (at 760 nm) of TO-PRO3 labeled nuclei in a thin section of RTF cleared mouse embryo. With a confocal aperture of 2 Airy, we found a factor of × 4,000 btween linear and non-linear excitation (2 μW vs. 8 mW/spot, respectively).

Based on these results, we decided to combine TO-PRO3 nuclear staining and RTF clearing for directly comparing the performance of confocal and OASIS microscopes.

### Faster and less invasive acquisition of 3-D image stacks

Tracing fluorescent dendrites and axons to study the projections and connectivity in small cellular net-works is a major goal of current neuroanatomy. At 250-nm lateral and micrometric axial sampling and with typical pixel dwell times of 1 (10 μs), sequential singlespot scanning schemes are necessarily slow, requiring 4 (40) ms, 4 (40) s and more than 1 (10) hrs for the acquisition of 3-D image stacks from cubes of 10 μm, 100 μm and 1 mm side-length, respectively. Parallelizing both the excitation and emission detection, as with our OASIS microscope is expected to considerably speed up the imaging of such large data sets.

Using TO-PRO3 nuclear staining as a proxy, we acquired *z*-stacks of images in RTF-cleared mouse embryos. Comparing the achievable imaging depths with the OASIS microscope in non-cleared (30 μm, **Fig. 3*A*** *and* **S5*A***) *vs.* RTF-cleared samples (90 μm, **Fig. 3*B*** and **S5*A***) we noted the 3-fold larger attainable imaging depth and well-preserved structural details. Next, we evaluated the photobleaching with the OASIS and confocal microscope. We first compared 2P-light sheet and confocal acquisition at shallow imaging depths, by continuously recording images of the same region of interest (ROI) at 30-μm depth in an non-cleared embryo. At the same initial signal-to-noise ratio for both microscopes, We observed a ∼3%-intensity loss after the first image with the OASIS microscope (*t*_exp_ = 480 ms per image, 15.7 mW/spot for the OASIS), whereas the signal remained relatively stable after a confocal scan. Fitting a monoexponential with the OASIS bleaching data revealed a 1/e (37%) loss of fluorescence every 21 frames, **Fig. 3*C***. Thus, fs-pulsed non-linear excitation results in a significantly higher pleaching in superficial tissue layers. However, for 3-D imaging of thicker sections, confocal microscopy rapidly produced much faster photobleaching than the OASIS because it required higher and higher laser powers to maintain image contrast at greater imaging depths; in fact, acquiring the first complete *z*-stack with the CSLM attenuated the TO-PRO-3 fluorescence so much that a second acquisition was impossible, **Fig. 3*D***.

**Fig. 3.**
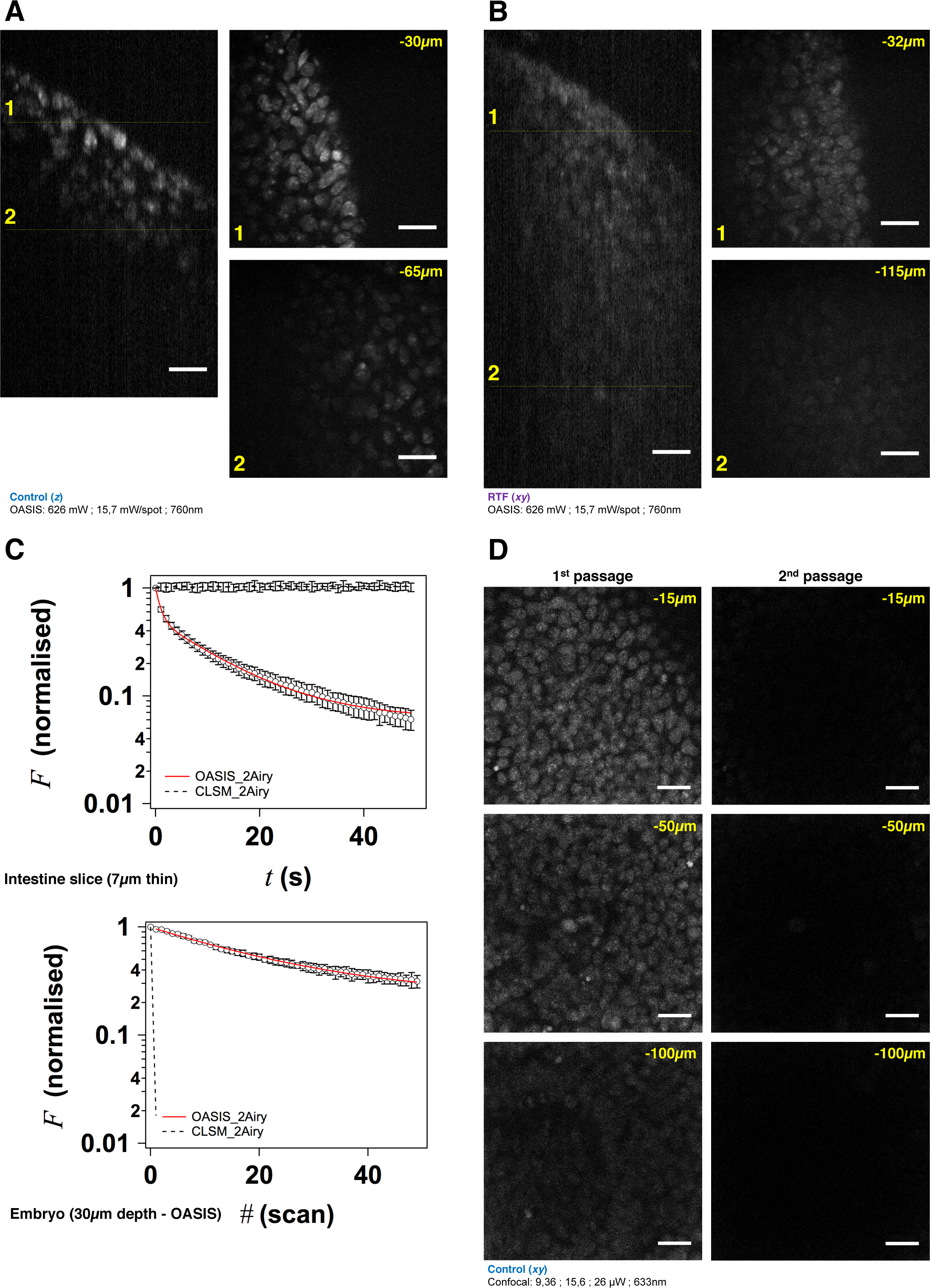
The OASIS microscope outperforms a CLSM for 3-D embryo imaging. **A.** 3-D data set taken with the OASIS in a non-cleared embryo stained with TO-PRO3 (Control). Panels show *xz*-projection of a *z*-stack of images (*right*) and *xy*-planes (*left*) corresponding to the dashed lines at 30 μm (1) and 65 μm imaging depth (2), respectively. Scale-bar, 25 μm. **B.** same, at 32 μm (1) and 115 μm depth (2), respectively, for an RTF-cleared embryo. Scale-bar, 25 μm, as in (A). **C.** Representative bleaching curves during continuous acquisitions, from a single *z*-section in – respectively -a thin slice of intestine (*top*) and of TO-PRO3-labeled nuclei (*bottom*) at 30-μm imaging depth in a embryo. **D.** Representative planes at various imaging depths (15, 50 and 100 μm, respectively) of a 3-D data set (200 planes from the surface to 200 μm, Az = 1qm;) acquired from a TO-PRO3-labled embryo. The 2 colums show the sections during the 1^st^ passage (*left*) and during the 2^nd^ passage, after completion of the 1^st^ image stack (*right*). Note the almost complete bleaching prohibiting repetitive volume imaging for the CSLM but not the OASIS microscope. Measured laser powers are given for each depth. Scale-bar, 25 μm.

We attribute the much higher volume photobleaching upon 1P confocal imaging in non-cleared embryos to four reasons, (*i*), tissue scattering at 633 nm was roughly double that of near-infrared light. The exponential scattering losses of excitation photons must be compensated for by exponentially increasing the excitation powers with increasing imaging depth; (*ii*), as a consequence of linear (1P) excitation, off-focus excitation of fluorophores located above and below the imaged plane causes bleaching, too, i.e., at any plane, bleaching occurs throughout the entire tissue volume while only one plane is imaged; (*iii*), although not contributing to imaging, the scattered 1P excitation light neverthelass excites (out-of-focus) fluorescence, which – in addition to the ballistic out-of-focus excitation – additionally contributes to photobleaching. Non-linear (2P) excitation, on the other hand, confines both fluorescence excitation and photobleaching to the focal plane, with the result fo better preserving the sample outside the plane which is actually imaged; (*iv*), image acquisition was 4-times faster on the OASIS compared to confocal scans (480 ms/image *vs.* 1.815 s/image for a similar image contrast), reducing the overall exposure of the sample.

We note that he better performance of the OASIS concept comes essentially from the excitation side, because with a confocal pinhole of 2 Airy, on the fluorescence collection side, both instruments should perform similarly. Finally, not only taking into account the loss of signal, but also that of Weber contrast, we found a three-fold larger effective depth penetration for the OASIS microcope, **Fig. S5*A***

Taken together, the OASIS microscope combines the advantages of 2P-exciation and wide-field imaging. Compared to the 1P-confocal, it achieves higher *z*-resolution, affords less photobleaching in 3-D samples and considerably speeds up data acquisition, thus allowing a more efficient and less invasive volume imaging.

### Fast volume acquisition from mouse embryonic bodies and brain organoids

We continued our comparison by imaging day-7 embryonic bodies (EB), **Fig. 4*A***, again stained with TO-PRO3. Images acquired at different depths displayed rounded structures with *lumina* inside tissue and revealed strong mitotic activity, **Fig. 4*B***. The rounded structures presumably correspond to neuroepithelial-like structures that are readily formed within EBs, indicating the inherent ability of the ectoderm to differentiate into neural lineages [42]. The sub-μm resolution of the OASIS microscope allowed us the detailed characterization of the different mitotic figures throughout the entire 160-μm thickness of the EB, **Fig. 4*C***.

**Fig. 4.**
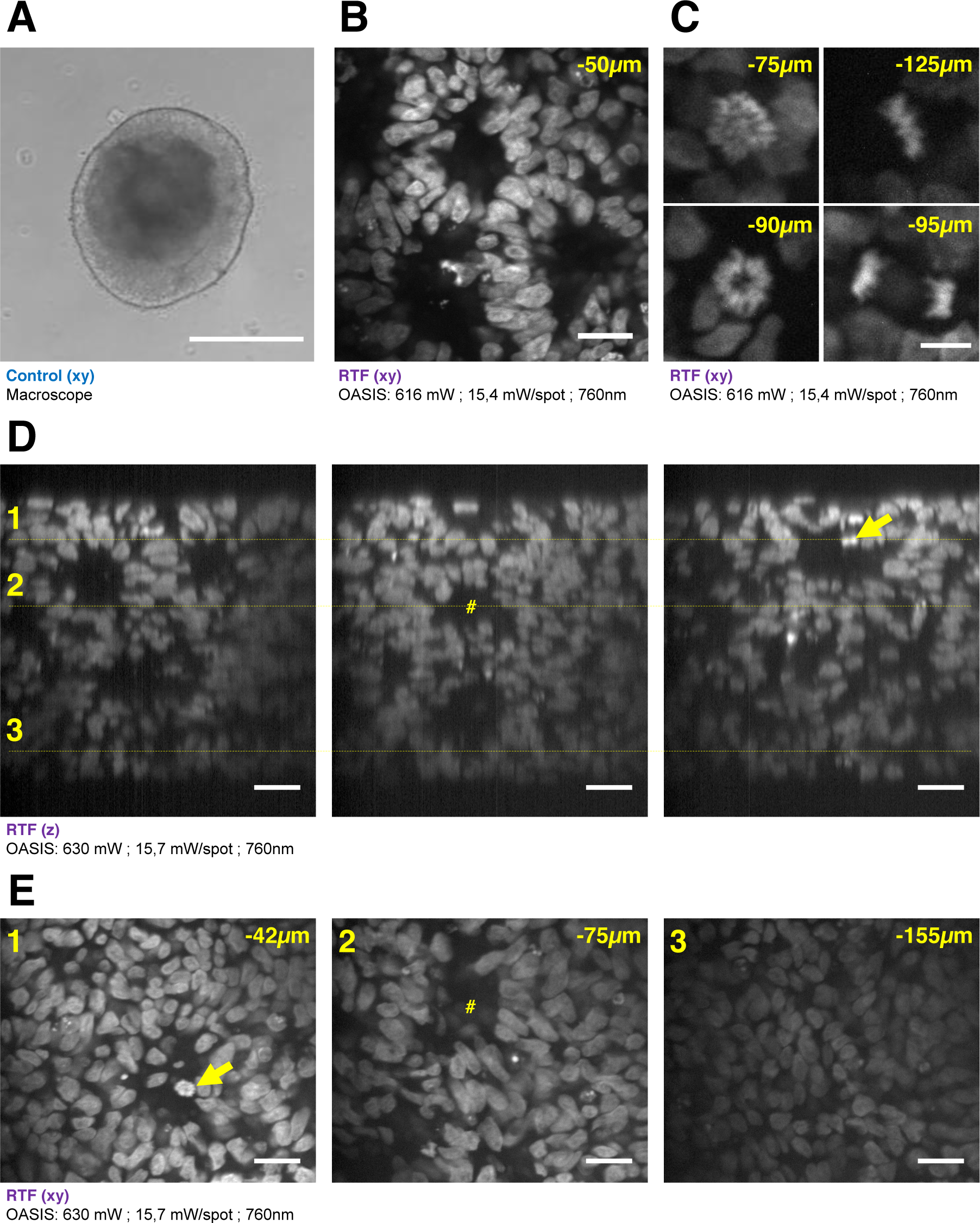
3-D imaging with the OASIS of the fine structure of embryonic bodies (EBs). **A.** Macrophotography of a day-7 EB; Scale-bar, 300 μm. **B.** Image acquired in the center of the organoid after TO-PRO3 staining, 50 μm below the surface. Scale-bar, 25 μm. Note the presence of internal round structures with a lumen. **C.** Zoom on mitotic figures observed in the EB at various depth (*top left*: prophase; *top right*: metaphase; *bottom left*: early anaphase; *bottom right:* late anaphase); stain: TO-PRO3. Scale-bar for all panels, 10 μm. **D.** *xz*-projections of a *z*-stack acquired across an entire day-7, TO-PRO3-stained EB. Scale-bar, 25 μm. Arrow points to mitosis also visible in panel (E). # indicates ventricle-like structure also perceived in (E). **E.** xy-sections across the dotted lines shown in (D) at imaging depths of, respectively, 42 μm (1), 75 μm (2) and 155 μm (3). Scale-bar, 25gm. With the exception of (A), all images were acquired after RTF clearing.

Acquirig a 3-D image stack at a *z*-spacing of 0.5 μm (**Fig. S5*B***) allowed us reconstructing complete EBs and realizing high-resolution projections along the orthogonal axes, **Fig. 4*D***, revealing fine structural detail and including again mitosis across its entire volume, **Fig. 4*E***. Volume imaging of entire EBs was almost 4-times faster with the OASIS compared to the CLSM (a 200-μm *z*-stack with a 0.5qm *z*-spacing required 3’12” vs. 12’6”), with no detectable photobleaching. Together, these features make the OASIS microscope an ideal setup for 3-D the detailed characterization of EB development.

Similar if not larger *z*-stacks were acquired a from RTF-cleared day-11 brain organoids, **Fig. 5*A***. At this early developmental stage, the neuroepithelium has been induced and forms buds that undergo 3-D growth within the Matrigel droplets [43]. Our observations highlight strong morphological modifications during this tissue expansion. A recurrent feature was that TO-PRO-3 labeled nuclei were rounded just below the surface of the brain organoids, whereas polymorph and diamond-shapes prevailed at greater imaging depths, **Fig. 5*B***. Also, the cell density and nuclear labeling changed markedly with depth. Orthogonal planes revealed compact groups of nuclei with stronger fluorescence, **Fig. 5*C***, as well as cavities and rounded structures with a neuroepithelium-like shape. As before in EBs, the resolution of the OASIS microscope allowed us to detect the presence of mitotic figures at the luminal side, **Fig. 5*D***. In brain organoids, typical achievable imaging depths were around 200 μm, reflecting the densification and opacification of the tissue during the development of an EB towards a brain organoid.

**Figure 5.**
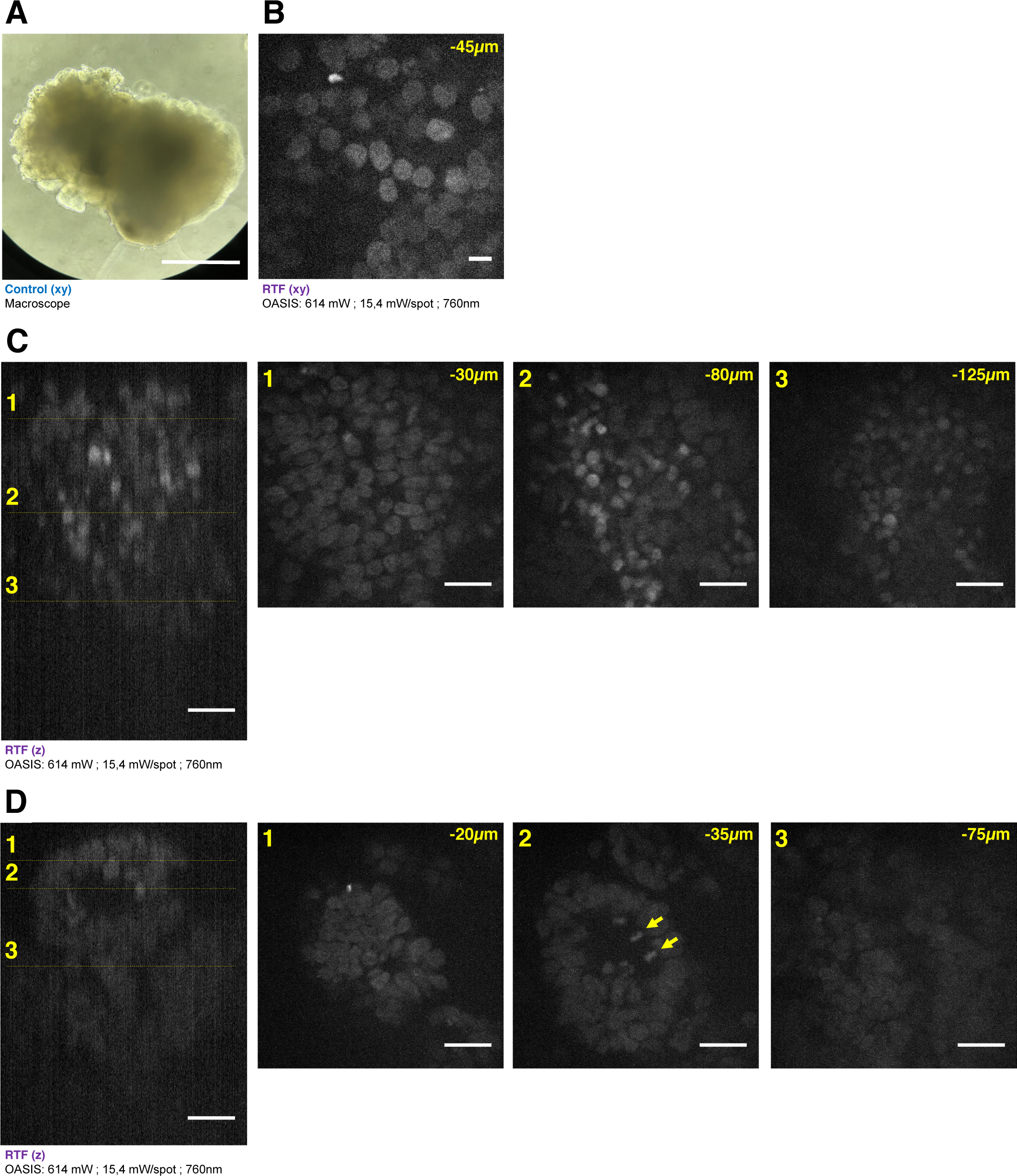
Brain organoid imaging with OASIS. **A.** Macrophotography of a day 11 brain organoid before clearing and observation on the OASIS microscope; scale-bar: 300 μm. **B.** Image acquired in the center of the organoid at 45 μm below the surface; stain: TO-PRO3; scale-bar: 10gm. **C.** *z*-reconstruction (left) and xy-acquisition (right) at respectively 30μm (1), 80μm (2) and 125μm (3) below the surface of the organoid; stain: TO-PRO3; scale-bar: 25μm. Note the change of the morphology of the nuclei in the center of the region of interest when depth increase. **D.** *z*-reconstruction (left) and *xy*-acquisition (right) at respectively 20μm (1), 35μm (2) and 75μm (3) below the surface of the organoid; stain: TO-PRO3; scale-bar: 25μm. Note the globular structure (neuro-epithelium) that can also be observed on the macroscopy and numerous mitosis inside this structure (arrows). Except in A. all images were acquired after the clearing of the organoid with the RTF protocol. Note the image quality across the entire imaged volume.

With its large field-of-view, increased depth penetration, low photobleaching and greater speed of acquisition compared to the CSLM, the OASIS optical scheme lends itself ideally to the observation of entire EBs and whole brain organoids. Thin sections can be studied at depth with sub-cellular (μm-) resolution, without mechanical slicing. The reduced complexity compared to a classical light-sheet microscope, its compact mono-block design and comparble ease-of-use make it an ideal companion for functional neuroanatomy. The ongoing integration of a compact, fixed-wavelength high-power fs-pulsed laser into this package will make the OASIS a unique, portable, alignment-free bench-top 2P-microcope.

## CONCLUSION

Work on brain organoids offers several distinctive advantages over classical disease models: (*i*), derived from patient fibroblasts, they raise less concerns than animal experimentation and work on human explants (see, however, [44] for an emerging awareness of the ethical issues associated with these 3-D cultures); (*ii*), they avoid the limitations of animal models that are often only a poor proxy of human pathology; (*iii*), they allow studies of rare or sporadic cases, for which genetic models are missing; (*iv*), they allow observing the onset of the disease during the early steps of the brain development, opening opportunities for studies that would be impractical or inacceptable on human embryos or infants.

For this field of applications, the OASIS 2P virtual light-sheet microscope is a compact, versatile and cost-efficient 2P wide-field research instrument allowing imaging of hiPSC cultures, embryonic bodies and brain organoids.

## MATERIALS AND METHODS

### Ethics statement

All experimental procedures were performed in accordance with the French legislation and in compliance with the European Community Council Directive of November 24, 1986 (86/609/EEC) for the care and use of laboratory animals. The used protocols were approved by the local ethics committee.

### Sample preparation

#### hiPSC culture and formation of embryonic bodies

Episomal human induced pluripotent stem cells (hIPSCs, Gibco) were cultivated on mitomycin-treated mouse embryonic fibroblasts using DMEM/F12 medium (Invitrogen), supplemented with 10% knockout serum (Gibco). When hIPSCs had reached about 80% confluence, they were detached with versene (ThermoFisher). Cell aggregates were removed and a single-cell suspension obtained with a cell strainer having a 100-μm mesh size (Corning). For the formation of embryoid bodies (EBs), 10^4^ cells were inoculated in 100 μl in each well of a ultra-low attachment, round-bottomed 96-well plate (Corning) and cultivated during 9 days in EB formation medium (StemCell Technologies).

#### Mouse embryos

Embryos were age E10.5 and E14.5. Mice were killed by cervical dislocation, the abdominal cavity was opened and the uterine horns were removed. Embryos were collected under a macroscope (Nikon SMZ800) and immersed in formalin (buffered 10% formaldehyde, VWR) overnight. They then were stored at 4°C in PBS / Sodium Azide 0,02%. E10.5 embryos were used for whole-embryo transparisation; E14.5 embryos were embedded in Optimal Cutting Temperature (OCT) compound and sliced into 7-μm-thin sections on a cryotome (Cryocut 1800, Leica).

### Staining, clearing and embedding

#### Nuclear staining

Samples were permeabilized by a 0.2% TritonX100 solution in PBS (during 15 min for 7-μm-thin embryo slices and EBs, 20 min for brain organoids, 1h for the whole-mount E10.5 embryo). They were then incubated overnight in a 1:1000 solution of TO-PRO3 (Invitrogen) or of chloroform-purified Methyl Green (Merck) in PBS and finally washed in PBS. At this point, non-cleared samples were mounted in a PBS-filled chamber under a glass coverslip for microscopy. We used a home-made recording chamber, that was modified from chambers designed for imaging and available as 3-D printer templates: https://idisco.info/idisco-protocol/).

#### Clearing

After nuclear staining, samples were processed for one of the three following clearing methods: TDE [39], Clear^T2^ [40] or RTF [41].

#### TDE

Samples were successively immersed in increasing concentrations (20%, 40% and 60%) of TDE (Sigma) solutions in PBS. The duration of each incubation varied as a function of the sample size: 1 h for 7-μm embryo slices and for EBs, 3 h for a whole E10.5 embryo.

#### Clear^T2^

Samples were immersed successively in, (*i*), a 25% formamide (Sigma) / 10% polyethylene glycol (PEG8000, Merck) solution in PBS (10 min for embryo slices and embryonic bodies, 30 min for a whole E10.5 embryo); (*ii*), a 50% formamide / 20% PEG8000 solution in PBS (5 min for slices and EBs, 15 min for whole embryo); (*iii*), a 50% formamide / 20% PEG8000 solution in PBS (1h for slices and for EBs, 3h for a whole E10.5 embryo).

#### RTF

Samples were successively immersed in, (*i*) a 30% triethanolamine (TEA, Sigma) / 40% formamide (Sigma) solution (15 min for slices and for embryonic bodies, 20 min for brain organoids, 3h20min for a whole E10.5 embryo); (*ii*) a 60% TEA / 25% formamide solution (25 min for slices and for embryonic bodies, 30 min for brain organoids, 5 h for embryo); (*iii*) a 70% TEA / 15% formamide solution (25 min for slices and for EBs, 30 min for brain organoids, 5 h for whole embryo). In either case, after clearing, samples were mounted under a glass coverslip in a chamber filled with the respective final solution.

### Microscopy

#### Confocal microscopy

We used a Zeiss LSM 710 microscope with a ×63/1.4NA oil-immersion objective, for the acquisition of the confocal data shown in figs. 2 and S3, and a ×40/1.0NA water-immersion objective for tissue sections and thick samples (embryo, EB, brain organoid), respectively. For a fair comparison of the performance for imaging thick samples of the confocal and 2P-virtual light sheet OASIS microscope, we set the confocal pinhole to 2 Airy units and the scanned area was restricted to 900 × 900 pixels with a zoom resulting in an effective pixel size of 0.182 μm, equivalent to the OASIS. We also acquired confocal images at 1 Airy for comparison. The laser powers delivered to the sample are indicated in each figure. For 1PEF, we excited TO-PRO3 using the 633-nm line of a HeNe gas laser (Lasos). Fluorescence was collected in between 646 nm and 725 nm.

#### 2P-virtual light-sheet microscopy

The optical path of OASIS microscope is schematized in Fig. 1. A detailed technical characterization is given in a companion paper, (Ricard, Deeg, *et al., submitted).* Briefly, 2P fluorescence is excited in 40 foci. These spots are scanned across the sample and the fluorescence emission from the spots confocally detected, similar to a Nipkow-Petráň spinning-disk microscope [45]. However, for optimal 2P excitation, a disk was manufactured that combines micro-lenses, dielectric long-pass (LP) coating and confocal pinholes. On the front side of this glass disk (borofloat B33, 100-mm diameter, 2-mm thick) >5,000 micro-lenses are arranged in four nested spirals were micro-machined (diameter *D* = 666 μm, focal length *f* = 7.8mm, lens-area-fill factor: 73.2%, the average inter-lens spacing is about 666 μm). The dielectric coating (T_ave_ < 0.05% for 400-675 nm, Tve > 93% for 730-1100 nm) on the rear side has pinholes (i.e. non-coted areas, each having 60-μm diameter) centered on the micro-lenses on the front side of the disk (see Fig. 1, *inset*).

For 2P excitation, the Gaussian beam of a fs-pulsed Ti:Saph laser (Spectra-Physics MaiTai HP with DeepSee™ module) was shuttered (AOM: AA Opoelectronic), expanded and collimated to a beam diameter of about 5 mm (1/*e*^2^) and aligned on the rear side of the disk. The disk transmits the IR light and the microlenses on the front side generate a spiral pattern of foci 7.8 mm in front of the disk. This pattern is imaged onto the sample by a telescope formed by the tube lens (*f* = 200 mm) and water-immersion objective (Nikon CFI75 Apochromat LWD 25× MP, NA 1.1w).

2PEF is detected through the same objective and tube lens. However, between the tube lens and the disk, two LP dichroics (custom design and manufacture from Alluxa, 50%-cut-on at 705 nm, T_ave_ < 1.5% from 400-695 nm, T_ave_ > 96% from 715 – 1100nm, flatness < 0.25 wave P-V per inch, *d* = 1mm) were used at 18° angle of incidence (AOI) to image the emission from the excitation spots onto the pinholes of the disk. As a consequence, the collected fluorescence from the excitation spots arrives focused at the level of the pinholes whereas out-of-focus fluorescence and scattered emission light is blocked. The intermediate image in the pinhole plane is separated from the IR excitation light by an AOI 45° primary dichroic LP filter (Custom design, manufactured by Alluxa, AOI 45°, flatness < 0.25 wave P-V per inch, *d* = 1 mm) and magnified by a. *f*_1_ = 140 mm and. f = 200 mm telescope onto an sCMOS camera (PCO.edge 4.2, chip size 13.3 mm × 13.3 mm). Due to this magnification, the image size on the chip is about 7.1 mm and the total magnification of the system is ×36. To block residual IR excitation light, a multi-photon-emitter (ET700SP-2P, Chroma) was used. For dual-color imaging a LP dichroic under AOI 15° (in fact a Semrock 532-nm laser BrightLine Di02-R532, AOI = 45°. For AOI = 15° its edge shifts to 568 nm) was used in combination with a knife-edge right-angle prism to split the image in the non-infinity space between the *f* = 200-mm lens and the camera chip. The red emission (>568nm) is imaged on one half of the sCMOS chip, and the green (<568nm) on the other, without further emission filtering. The fluorescence signal at the margin of the field of view is relatively weak, and cropping the image of about 7.1-mm diameter to the half chip size of 6.65 mm x 13.3 mm is justified. Image cropping, rotation and alignment were automatized in a custom-written macro in FIJI [46].

The upright microscope body was constructed as a rugged monocoque, machined from a single metal block (see photos), with slots for the optical and opto-mechanical elements at pre-defined places. A highly folded realization of the beam-path optimized by ZEMAX allowed the compact design. We used a voice-coil based *z*-objective drive, combining accuracy and precision over a wide focusing range. A flipable mirror was used to toggle between 2P-imaging and trans- or reflected-light imaging. This alternative beam path uses a different *f* = 140-mm tube lens in combination with a small USB camera (Point Grey, BFLY-U3-23S6M-C). Having an effective magnification of ×17.5 this imaging mode was used for getting an overview of the sample and for identifying regions of interest (ROIs) to be imaged in 2P fluorescence. All image acquisition parameters, the laser and the objective focus drive were controlled through an in-house microcontroller imaging software. The used excitation wavelength λ_ex_, laser power *<P>*, and *z*-spacing Δ*z* for stacks were software controlled and are specified in the figure legends. Images were acquired without software binning at the full sCMOS resolution.

### Image analysis and quantification

Images were analyzed and displayed using FIJI. For a better visibility of the faint fluorescent signals at greater imaging depths, we used a nonlinear look-up table (*γ* = 0.6) for Figs. 4B-E and 5C-D.

Weber contrast was calculated as C_w_ = (*F-B*)/*B*, where *F* and *B* are the fluorescence intensities of the image and background, respectively. *B* was measured as the mean intensity of a cell-free region and *F* corresponded to the mean intensity in a small ROI (18 μm x 18μm) in the center of the image for fig. S5A, and to the whole image in fig. S5B. (In fig. S5A the sample was inclined respect to the optical axis so that only a small region could be used to quantify the penetration depth).

### Statistics

Results are at least triplicates of three experiments and are represented as mean ± SD. Student’s t-test was used to compare among experiments. Data were processed and figures prepared using IGOR Pro (Wavemetrics).

## Footnotes

1 The ‘*brain in a vat*’ or ‘*brain in a jar*’ is a is a scenario used in a variety of *Gedankenexperiments* intended to draw out certain features of human conceptions of knowledge, reality, truth, mind, consciousness and meaning. It is an updated version of René Descartes’ evil demon thought experiment. It has been extensively used by Hilary Putnam (*Reason, Truth and History*, 1981), in an argument inspired from Roald Dahl’s short story, « William and Mary » (1959).

## List of abbreviations

2P: two-photon
3-D: three-dimensional
AOI: angle of incidence
BSA: bovine serum albumin
CLSM: confocal laser scanning microscope
DMEM: Dulbecco’s modified Eagle medium
EB: embryonic body
FWHM: full-width at half maximumhi
PSC: human inducible pluripotent stem cell
LP: long-pass (filter)
MG: Magnesium Green
NA: numerical aperture
OASIS: <u>O</u>n-<u>a</u>xis 2-photon light-<u>s</u>heet generation *in-vivo* <u>i</u>maging <u>s</u>ystem
OCT: opimal cutting temperature
PBS: phosphate-buffered solution
PEG: polyethylene glycol
RI: refractive index
ROI: region of interest
RTF: <u>R</u>apid clearing method based on <u>T</u>riethanolamine and <u>F</u>ormamide
sCMOS: scientific Complementary Metal Oxide Semiconductor
SD: standard deviation
SPIM: selective-plane illumination
TDE: 2,2’-thiodi ethanol
TO-PRO-3: a carbocyanine monomer nucleic acid stain with red excitation and far-red fluorescence (642 nm/661 nm) similar to Alexa Fluor 647 or Cy5. It is among the highest-sensitivity probes for nucleic acid detection.

## Author contributions

AD and RU designed and conceptualized the OASIS prototype, BD and PD generated iPSCs, embryonic bodies and brain organoids, AD, BD, IR, CR, MB and MO performed experiments, MO wrote the paper with contributions from all authors.

## Acknowledgements

The authors thank Dr Elke Schmidt for help in setting up early experiments, Patrice Jegouzo (fine mechanics and 3-D printing) as well as Claire Mader and her team (CNRS FR3636 Saints Péres Central Animal Facility) for expert technical support. Confocal imaging was done at the SCM core facility (*Service Commun de Microscopie,* with the support of Jennifer Corridon, Paris Descartes). We thank Dr. Christian Seebacher (TILL.id) for ZEMAXing and optimizing the optical design.

Financed by the European Union (H2020 Eureka! EUROSTARS project ‘OASIS’, to RU and MO), the CNRS (DEFI *Instrumentation aux Limites,* to MO), the *Agence Nationale de la Recherche* (ANR-10-INSB-04-01, *grands investissements* France-BioImaging, FBI, to MO) and the Region Ile-de-France (DIM *canceropole,* project EDISON, to MO).

The Oheim lab is a member of the C’nano IdF and *Ecole de Neurosciences de Paris* (ENP) excellence clusters for nanobiotechnology and neurosciences, respectively.

